# Raven Calls Indicate Sender’s Neural State

**DOI:** 10.1101/613778

**Authors:** Zixuan Huang, Zhilong Wang, Jun Xie, Greg Mirt, Chengying Yan, Jing Zhong, Xianli Deng, Fangfang Liu, Chunlin Zhou, Fan Xu

**Author notes:** These authors contributed equally (Zixuan Huang, Zhilong Wang, Jun Xie, Greg Mirt). Co-corresponding Authors: Prof Chunlin Zhou, School of Geography Science, Nanjing Normal University, Nanjing, Jiangsu Province, CHINA, 210023, Prof Fan Xu, Department of Public Health, Chengdu Medical College, Chengdu, Sichuan Province, CHINA, 610500.

## Abstract

Vocal communication accounts for dominantly percentage within animal species. The information of vocal samples contains not only the amplitude of objects, but also the emotional states behind it. However, to extract the emotion state behind the sound remains controversial. Here we introduce an artificial network method, the Back Propagation Neural Network, BPNN, to classify the emotional states behind the sound. The results disclosed the behaviour categories, including alarm, flight, begging and singing which has been successfully classified. This artificial intelligence classification may aid us to distinguish the ecological categories via animal vocal communication and to discover its significance of evolution and nature.

## Background

The sound is one of universal phenomenon, which does not only exist in numerous nature phenomenon [1–3], but also appear in animal communication in species or across species [4, 5]. From the perspective of physics, the sound is generated by the amplitude of object, which determines the frequency and intensity [6–8]. According to evolution, the ability of adaption and survival skills of species had been constructed and strengthened for surviving, for example: track the life source, avoid the harmful events and defend the territory [9, 10].

With respecting the animal communication, there are three domain categories, sound, behaviour and the order [11–14]. The pronunciation process of animal depends on a series of muscles interaction, including breathing muscles, mouth muscles and throat muscles and the amplitude of vocal cord [15, 16]. Vocal communication accounts for dominant percentage in species [17, 18]. Sound response on the stimulation is the best indicator of sender’s emotional state under the specific circumstances, such as nervousness, sadness, happiness, repulsiveness, angriness and afraid [19, 20].

Interestingly, the raven, an intelligent avian, has been used in many specific mind and brain studies for decades [21–23]. For example, the raven’s memory predict future [24], sequence tool used to get the award [25], the awareness of density of article[26], and so on. Ravens’ call often appears frequently during their foraging[27], defend the territory[28], rutting and mating[29], infant parenting[30]. Even though we can hear various different callings, however, to classify the emotional states behind the sound remains unclear. Here are four behaviour validated related calls of raven are hypothesized to be directly different among acoustic parameters, providing cues to emotional category recognition. Therefore, we use the method, termed as “Back Propagation Neural Network, BPNN”, to disclose the emotion behind the sound. This may be helpful to us to understand each other better, not only in same species, but also across species.

## Method

### Data Source

Total 851 sound samples of raven have been downloaded from the website (https://www.xeno-canto.org/species/Corvus-corax) and their detail information sorted into excel file, totally 11 variables included: “Common name, scientific name, recording duration, recording person, date and time of recording, country, location, altitude, tweet type, behaviour marker, audio file classification number”. The variable “type” described the behaviour profiles according to the current situation, including alarm, fighting, flying, begging, and singing. Furthermore, the “Remarks” listed the detail raven’s behaviours. Aim to reduce the noise impact and enhance the discrimination effect, the sound samples, which contain more than 2 raven’s sounds, have been excluded. Four different categories of sound samples (alarm, singing, fighting and begging) have been selected. Consequently, there were only 100 sound samples recruited in the final data analysis.

### Acoustic Parameters Extraction

Here we use the MATLAB script to extract the acoustic parameters [31, 32]. Including: (1) Mean fo(Hz), (2) Maximum fo(Hz), (3) Minimum fo(Hz), (4) Range fo(Hz), (5) Start fo(Hz), (6) End fo(Hz), (7) fo at the half of call duration (mid fo Hz), (8) slope from fo start of the call to the fo maximum (slope S-M; Hz/s), (9) Slope from maximum fo to the end of the call (Slope M-E; Hz/s), (10) Inflection rate (number of frequency changes/s), (11) Harmonicity (HNR, dB), (12) Jitter (the absolute; fo difference between consecutive fo measurements/ the average period), (13) fo variation (sum of all fo changes measured/ call duration, Hz), (14) amplitude range (maximum dB-minimum dB).

### Data Analysis

According to the raven’s call recorded under the different circumstances, the biological meanings may be presented differently. Therefore, we selected Back Propagation Neural Network (BPNN) method [33–35]. A typical BPNN consists of the input layer, hidden layer and output layer (Figure 1).

**Figure 1.**
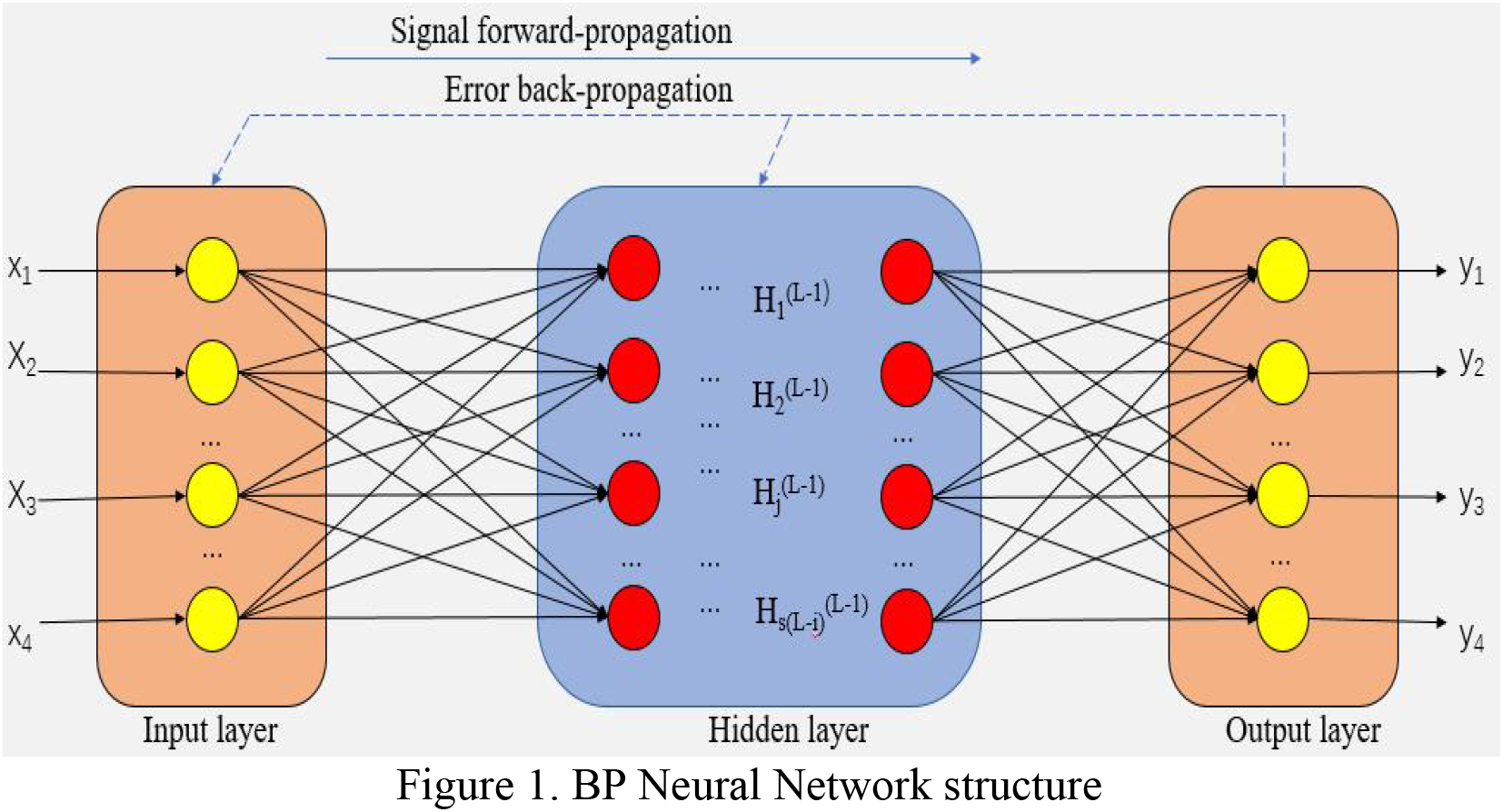
BP Neural Network structure

## Result

Totally we have selected four different categories call of raven, including alarm, begging, fighting and singing. The descriptive acoustic parameters extracted from MATLAB, please see Table 1.

**Table 1.** Four different Call Sound Parameters

| Parameters | ALARM |  |  | FLIGHT |  |  | BEGGING |  |  | SONG |  |  |
| --- | --- | --- | --- | --- | --- | --- | --- | --- | --- | --- | --- | --- |
|  | N | means | s.dev. | N | means | s.dev. | N | means | s.dev. | N | means | s.dev. |
| Amplitude range | 10 | 0.7107576 | 0.5005164 | 74 | 1.0111211 | 0.572916 | 6 | 0.9334848 | 0.746398 | 10 | 0.894717 | 0.624794 |
| *fo* variation | 10 | -2.676767 | 6.7065086 | 74 | -139.71842 | 1094.613 | 6 | -8.852867 | 18.1973 | 10 | -68.3103 | 144.6527 |
| Start *fo* | 10 | -1.83E-05 | 3.86E-05 | 74 | 0.0002924 | 0.002695 | 6 | -2.035E-05 | 4.98E-05 | 10 | -8.2E-05 | 0.000261 |
| Harmonicity | 10 | -21.22948 | 7.2070505 | 74 | -21.673169 | 5.745358 | 6 | -25.54435 | 8.567675 | 10 | -19.8766 | 6.758902 |
| Inflection rate | 10 | 2.46E-09 | 5.63E-09 | 74 | -4.60E-10 | 1.78E-08 | 6 | -1.37E-09 | 3.76E-09 | 10 | 1.50E-08 | 4.78E-08 |
| Jitter | 10 | 2.81E-09 | 6.22E-09 | 74 | -1.40E-09 | 1.06E-08 | 6 | -1.16E-09 | 4.10E-09 | 10 | 1.07E-08 | 3.35E-08 |
| End *fo* | 10 | 0.0044069 | 0.0105198 | 74 | -0.001516 | 0.010735 | 6 | -0.001943 | 0.0112 | 10 | 0.001276 | 0.003853 |
| Maximum *fo* | 10 | 0.3562669 | 0.2590624 | 74 | 0.4943541 | 0.284507 | 6 | 0.4442582 | 0.350765 | 10 | 0.437178 | 0.31567 |
| Mean *fo* | 10 | -6.41E-06 | 2.284E-05 | 74 | -0.0007544 | 0.006447 | 6 | -1.181E-05 | 1.73E-05 | 10 | -0.00165 | 0.003674 |
| Middle *fo* | 10 | -0.000305 | 0.0200354 | 74 | -0.0023711 | 0.037413 | 6 | 0.0065875 | 0.017132 | 10 | -0.00995 | 0.050983 |
| Minimum *fo* | 10 | -0.354492 | 0.2501229 | 74 | -0.5167672 | 0.293957 | 6 | -0.4892283 | 0.405914 | 10 | -0.45754 | 0.312752 |
| Range of *fo* | 10 | 0.7107576 | 0.5005164 | 74 | 1.0111211 | 0.572916 | 6 | 0.9334848 | 0.746398 | 10 | 0.894717 | 0.624794 |
| Slope M-E# | 10 | -2.27E-06 | 5.011E-06 | 74 | -1.191E-06 | 1.87E-06 | 6 | -7.04E-07 | 1.43E-06 | 10 | -3.9E-06 | 4.69E-06 |
| Slope S-M | 10 | 3.72E-07 | 2.99E-07 | 74 | 3.493E-06 | 6.68E-06 | 6 | 1.131E-06 | 1.4E-06 | 10 | 3.23E-06 | 4.69E-06 |

By meaning of BPNN method, the biological meanings of raven’s call have been classified. Firstly, we have selected the training samples: 14 sound parameters were used. We tested first 100 sound samples. Namely, all data imported training dataset. Secondly, pre-processing: Due to realistic complexity, we used the standard process of deviation normalization (x. std. = (x-min) / (max-min)) to perform the better neural network and to reduce the negative effect of over fit. The new data was constructed via dimensionless processing. Thirdly, primary parameter selection: The number of input layer units was 14. There was only one output layer neuron. Then, the number of hidden layer neurons was 12. Details please see Figure 2.

**Figure 2.**
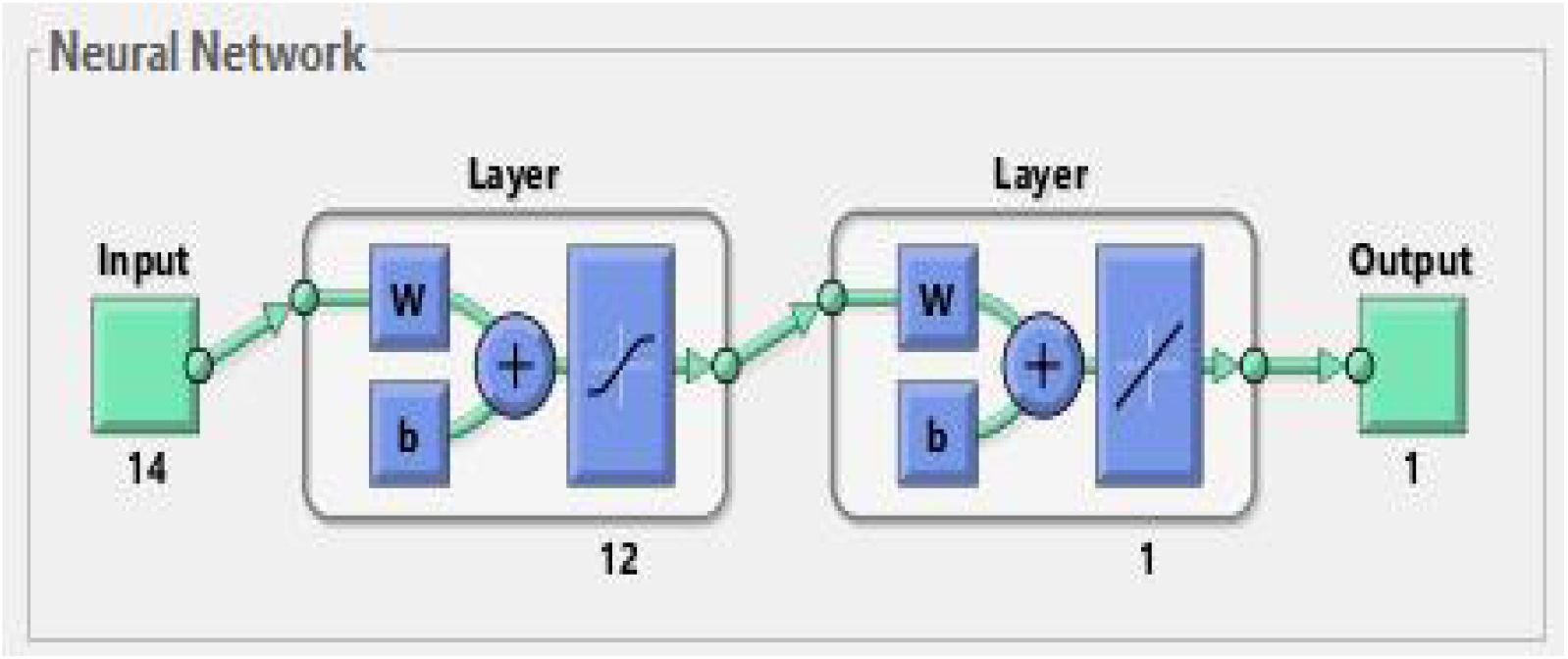
Neural Network Pattern

The unique purpose of BPNN construction is to classify. Double transfer sigmoid function and linear were used. The results were presented well when used “tansig” function in the input layer-hidden layer and linear function in the hidden layer-output layer. Furthermore, the “trainlm” method (combined gradient descent method with Newton method) was used for training, which terminated when achieved the best effect. The details of training performance process please see Figure 3.

**Figure 3.**
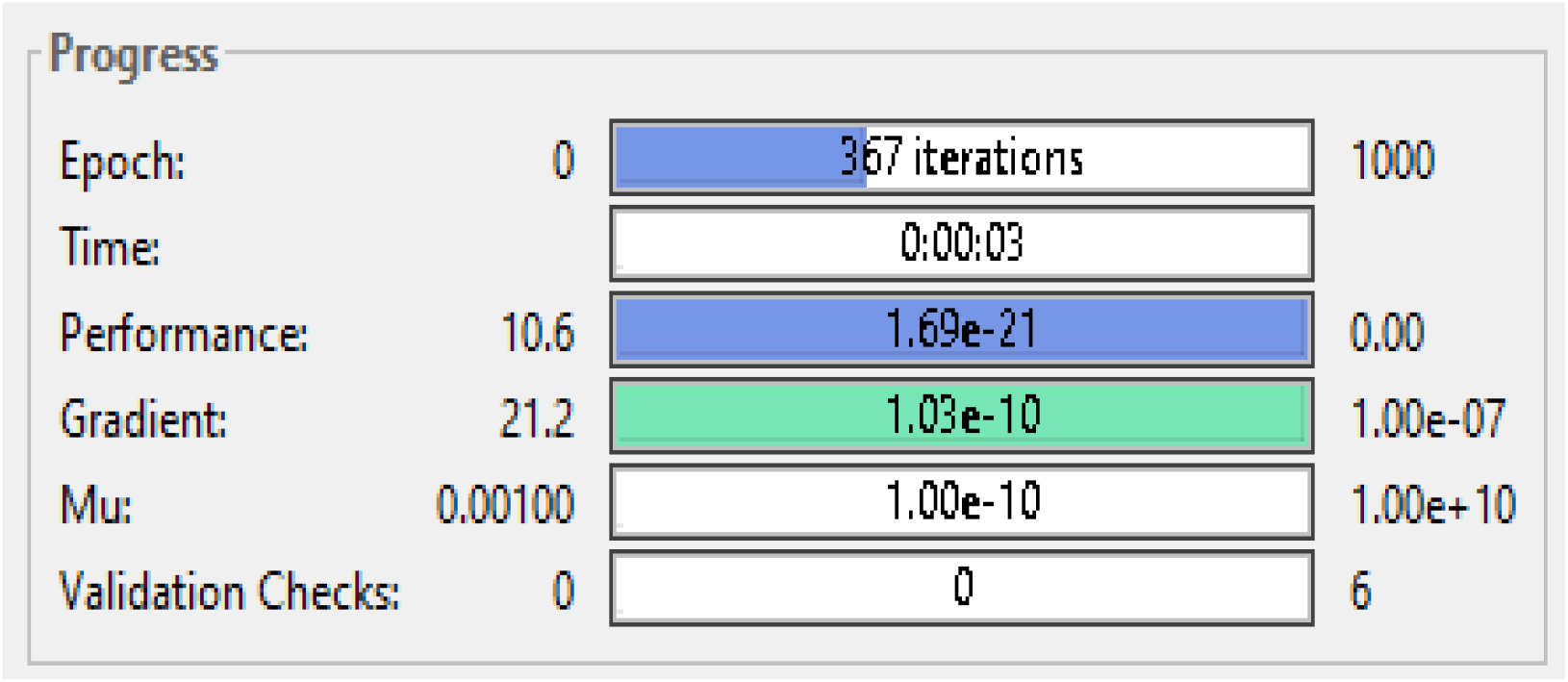
Training Process

Fourth, Validation the prediction effect: the training network was used to predict the types of raven calls, to detect the prediction and promotion ability of the network, and to get the classification of network simulation and actual classification. The detail comparison results please see figure 4.

**Figure 4.**
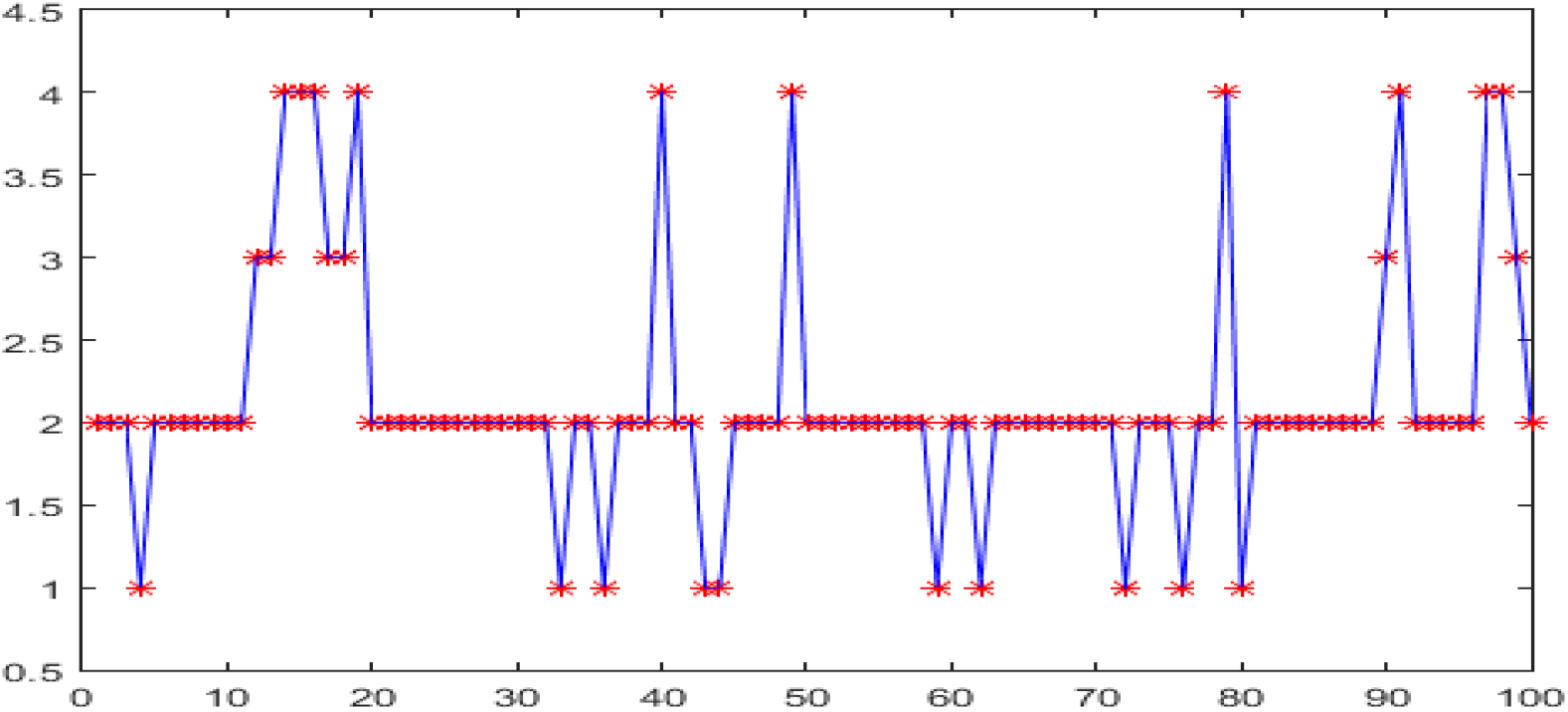
Prediction effect Figure legend: red star stands for actual classfication, blue line stands for stimulation classfication

The overall trend of the predicted value was close to the actual value. It can be inferred that the prediction of the model is more accurate, while it has a certain reference value. The results disclosed that the average absolute error and the average relative error of the actual classification and the predicted classification were 1.0e-09 and 1.0e-10, respectively.

## Discussion

The sound is the best indicator to reflect the property of the object [36, 37]. From the physical perspective, the size of sound depends on the frequency of object, and the pitch of sound depends on the amplitude of object [38, 39].

With respect to the animal communication, the pronunciation is not only depending on a series of pronounced organ coordination work [40–42], but also depending on the control of central nervous system [43]. The animals have been adapted to the environment after a long time evolution process via a series of successful constructed conditional reflexes [44]. It is obvious that the emotional information maybe hide behind the vocal communication.

The raven is recognized as the most intelligent avian [45, 46]. It was being used in mind, sequenced tool used test, awareness, learning and memory researches. More interestingly, Markus et al distinguished the gender and age of raven [47], and reported that ravens can predict future via their memory [24].

There are four behaviour categories of raven’s call, including alarm, begging, flighting and singing. It is necessary to distinguish the specific behaviouristic category. Thanks to BPNN method, their behaviour categories have been classified successfully.

The abundance information is contained within the sound in animal communication. There is not only the amplitude information of object, but also embedded emotional state. For example, professional police can distinguish the criminal suspects via sophisticated conservation due to their unstable irregular speaking [48]; experienced hunter can determine the degree of hunger of animal [49].

To distinguish the emotional state behind the sound is of great significance. Through the emotion information behind the ravens’ sound, we can get better understanding of their biological meanings. For instance, the alarm call often occurs on the occasion of dangerous events approaching, it is good for avoiding the harmful events and surviving [50, 51]. The flight call occurs on the situation of fighting process; the failure subject presents submission to dominant subject to avoid the further damage on its body or relationship [52, 53]. Moreover, the begging call happens when sub-dominant subject paying / begging for the life sources, such as the food, resting place etc [54, 55]. As for the singing; this may arise on the situation of pleasant animal communication, as for the specific motivation remains controversial.

Taken together, this artificial network could be helpful of trying to distinguish the behaviour categories of different raven’s sounds. It can strengthen our understanding on their biological and evolution meanings.

## Conclusion

Taken together, we herewith disclose the method of BPNN could be the promising candidate to classify the category of raven’s call. Furthermore, this may help us to investigate more in-depth of neurobiology of raven’s mind.

## Abbreviations List

BPNN: Back Propagation Neural Network

## Declarations

### Ethic Statement

Not applicable.

### Consent for publication

Not applicable

### Availability of Data and material

All data will be open for reasonable request.

### Authors’ Contribution

X.F drafted the general idea and drafted the manuscript with HZX; XJ, ZJ, YCY and DXL collected the sound data from xeno-canto website and data clean.HZX and WZL performed the MATLAB analysis.

## Acknowledgement

We thanks Dr Huang Kun provided the valuable data mining strategy and Peter proofreading and ensures the general language quality.

